# A multiplex real-time PCR assay for detection of equine herpesvirus 1 and equine herpesvirus 4

**DOI:** 10.1101/2025.03.31.646299

**Authors:** Rebecca L. Tallmadge, Melissa Laverack, Beate Crossley, Diego G. Diel

## Abstract

Equine herpesvirus (EHV) 1 and EHV-4 are common viral pathogens of horses that can cause upper respiratory disease, neurological disease, abortion, or death. As characteristic alphaherpesviruses, both EHV-1 and EHV-4 can establish latency, resulting in a lifelong carrier state in infected animals. Here we describe the development and validation of a rapid and sensitive multiplex real-time PCR assay (EHV1-4MP) that simultaneously detects EHV-1 and EHV-4 and includes an endogenous internal control targeting the equid genome. The EHV1-4MP assay analytical sensitivity was determined to be 15 genome copies for EHV-1, EHV-4, and equid MC1R per reaction. Analytical specificity was determined using a panel of 28 equine respiratory pathogens and commensal equine microorganisms. The EHV1-4MP assay detected reference and clinical isolates of EHV-1 and EHV-4, and did not detect other equine herpesviruses such as EHV-2, EHV-3, EHV-5, or several other viral and bacterial pathogens of horses. Importantly, the EHV1-4MP assay developed here has improved specificity compared to existing assays and is able to exclude the closely related EHV-3, EHV-8, and −9 viruses. Diagnostic performance was evaluated using 60 clinical samples including upper respiratory swabs and washes, blood, placenta, lung, and brain. The EHV1-4MP assay results were in 100% concordance with singleplex EHV-1 and EHV-4 assays. Our results demonstrate that the EHV1-4MP real-time assay developed here offers rapid, sensitive, and simultaneous detection of EHV-1 and EHV-4.

**Importance:** Equine herpesvirus (EHV) 1 and −4 are highly contagious and ubiquitous pathogens that commonly cause fever and respiratory disease, and can lead to outbreaks, abortion, neurological disease, or death. Here we describe the development and validation of a rapid and sensitive multiplex real-time PCR assay (EHV1-4MP) that simultaneously detects EHV-1 and EHV-4 as well as an endogenous equid control and an exogenous DNA control, with improved specificity compared to existing assays.

## INTRODUCTION

Equine herpesvirus (EHV) 1 and −4 are highly contagious and ubiquitous pathogens that commonly cause fever and respiratory disease. EHV-1 can achieve systemic distribution and in severe cases, results in abortion or neurological disease known as equine herpesviral myeloencephalopathy (EHM), while EHV-4 infection and symptoms are usually limited to the upper respiratory tract (1, 2). Generally, EHV-1 causes more severe disease than EHV-4, although the prevalence of EHV-4 is higher (3–5). In recent quantitative PCR-based surveillance studies of horses presenting with fever and/or respiratory signs, EHV-1 was detected in 0.7 to 3.0% of the cases, contrasting with EHV-4 which was detected in 6.6 to 10.8% of cases (6–8). Retrospective studies of infectious abortions in horses revealed that EHV-1 caused 3-6% of infectious abortions in Australia, United Kingdom, and United States, and up to 26% in Poland (9–12), whereas EHV-4 was associated with only 0.3% of the abortions in the United Kingdom (11). Outbreaks of encephalomyelitis and/or abortion in horses due to EHV-1 infection have been reported in the United States, Europe, Ethiopia, New Zealand, and China (13–20). A few outbreaks have been attributed to EHV-4; those that have been reported describe respiratory disease of horses and donkeys in Denmark, Germany, and Romania (21–23).

EHV-1 and −4 are members of the subfamily *Alphaherpesvirinae* and genus *Varicellovirus*, along with EHV-3, EHV-6, EHV-8, and EHV-9 (24). EHV-3, also known as equine coital exanthema, is distinct from EHV-1 and EHV-4 in that it causes mucocutaneous disease and is spread through venereal transmission rather than causing respiratory or neurological symptoms or abortion (25). EHV-6, −8, and −9 were not initially isolated from horses (24). EHV-6 was originally named asinine herpes virus (AHV) 1 because it was the first herpesvirus isolated from a donkey (26, 27). EHV-8 (AHV-3) was initially isolated from a donkey presenting respiratory symptoms; additional cases of respiratory disease, viral encephalitis, or abortion have been reported in donkeys and horses (26–31). Notably, two EHV-8 abortion cases in horses were misdiagnosed as EHV-1 due to PCR cross-reactivity (31). EHV-9 was isolated from a captive gazelle and hence named gazelle herpesvirus 1, subsequently it has been shown to be pathogenic in horses (32, 33).

EHV-1 infection of non-equid hosts including polar bears, black bears, Thomson’s gazelles, and guinea pigs in zoo settings manifested as neurological disease and often resulted in death of affected animals (34). Occasionally EHV species recombine with each other or with “zebra EHV-1”, which can lead to disease and death in unexpected species (35, 36).

Control of EHV-1 and EHV-4 is difficult in part because foals can be infected by two weeks of age, regardless of vaccination protocols (37). Also, EHV-1 and EHV-4 establish latency after infection, consistent with other alphaherpesviruses, creating an invisible reservoir of virus that can reactivate from latency and transmit to other susceptible animals (38–40). Notably, a study of 70 routine necropsies following musculoskeletal injuries identified latent EHV-1 infection in 25.7% of horses and latent EHV-4 in 82.8% (41).

Clinical symptoms are shared among respiratory infections caused by equine viral and bacterial pathogens, necessitating specific and sensitive diagnostic assays (7). Rapid diagnosis of EHV-1 and EHV-4 is imperative considering their highly infectious nature and potential for severe disease outbreaks. Real-time PCR-based diagnostics are the recommended method for direct detection of EHV-1 infection when respiratory symptoms are present and to manage quarantines and biosecurity during outbreaks (13, 42, 43). Likewise, the World Organization for Animal Health (founded as OIE) identified PCR as the standard method of EHV-1 and EHV-4 detection followed by virus isolation (44).

Here we describe a multiplex real-time PCR assay that was developed and validated to simultaneously test for EHV-1 and EHV-4. The built-in eqMC1R assay provides an endogenous extraction control to detect equid DNA in samples and verify sample integrity. An exogenous extraction control is also included as a means of monitoring assay performance.

## MATERIALS AND METHODS

### Real-time PCR assay design

D*e novo* real-time PCR assays targeting the EHV-1 glycoprotein B (gB) gene ORF33, EHV-4 gB ORF33, and the equid MC1R gene (Table 1) were designed with Primer3 web service (45, https://bioinfo.ut.ee/primer3/) and Primer3 2.3.7 in Geneious Prime 2023.1.2 to enable specificity testing (Biomatters, Inc., Boston, MA). All 135 EHV-1 and 50 EHV-4 sequences available in the NCBI nucleotide database at the time of assay design, as well as 12 EHV-1 sequences determined at the Virology Laboratory at the Animal Health Diagnostic Center, were used to build sequence alignments and identify regions with high sequence conservation. Representative gB sequences from EHV-1, EHV-2, EHV-3, EHV-4, EHV-5, EHV-6, EHV-8, and EHV-9 (NC_001491, HQ247739, NC_024771, NC_001844, AF050671, MT012704, MF431611 and MW822570, and NC_011644, respectively) were analyzed to confirm optimal specificity.

**Table 1.**
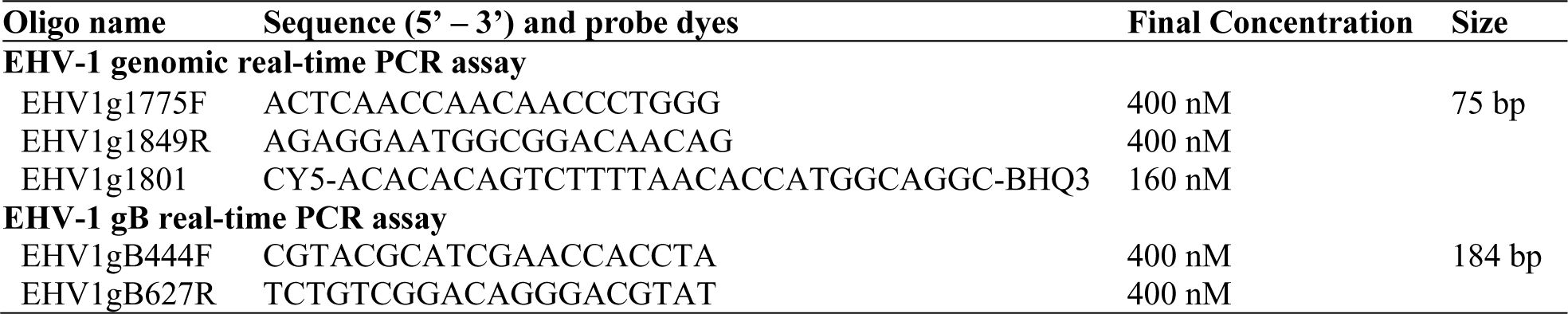
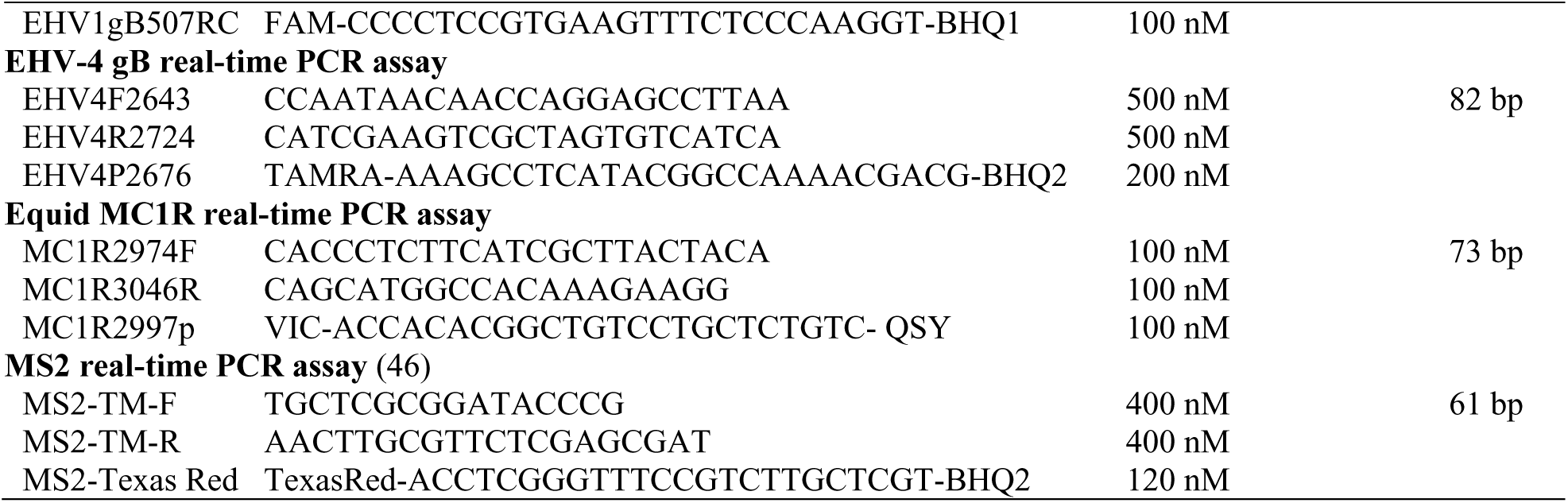
Equine herpesvirus 1 and 4 multiplex real-time PCR assay primer and probe sequences.

MC1R sequences were obtained from the NCBI nucleotide database on June 3, 2022 from the following equids and aligned to identify conserved regions: horse (*Equus caballus,* n=21), Przewalski’s horse (*Equus przewalskii*, n=2), donkey (*Equus asinus,* n=6), kiang (*Equus kiang,* n=2), onager (*Equus hemionus,* n=3), and zebra (*Equus burchelli, Equus grevyi, Equus zebra,* n=5). The forward and reverse primers and the probe are identical to these 39 sequences except for one nucleotide mismatch in the probe sequence compared to the 2 *Equus grevyi* sequences at position 7 out of 24 from the 5’ end. The equid MC1R real-time PCR assay primers and probe are within a single exon of the MC1R gene which enables detection of either genomic DNA or mRNA. In addition to the MC1R assay, an exogenous extraction control real-time PCR assay targeting the *Escherichia coli* bacteriophage MS2 was also included our multiplex assay (46).

Primers were synthesized by Thermo Fisher Scientific, Inc., Waltham, MA. Hydrolysis probe dyes were selected to facilitate multiplexing (synthesized by Sigma-Aldrich, St. Louis, MO, or Thermo Fisher Scientific).

### Viral and bacterial isolates, positive amplification controls, and clinical samples

Well-characterized EHV-1 and EHV-4 isolates (VLS 1246, VLS 1397) that are used in routine diagnostic testing and have been confirmed by whole genome sequencing were used to characterize assay performance. A panel of characterized equine respiratory viral (n=16) and bacterial (n=12) isolates were used to evaluate assay specificity (Supplemental Table 1).

Synthetic positive amplification controls (PACs) were designed for each assay with the target sequence region centered among 395-400 base pairs (gBlocks, Integrated DNA Technologies, Coralville, IA). Synthetic fragments of EHV-8 and EHV-9 gB regions were also obtained, based on NC_075566 and NC_011644 sequences.

A plasmid containing 200 bases of the MS2 TM2 region from GenBank nucleotide sequence V00642 was synthesized and cloned into pUC57 by GenScript USA, Inc (Piscataway NJ) to serve as an exogenous control.

Residual clinical specimens submitted for diagnostic testing at the Animal Health Diagnostic Center (AHDC) of Cornell University between January 18, 2019 and April 4, 2023 and archived at −80°C were used to evaluate the diagnostic sensitivity and specificity of the EHV1-4MP assay. A panel of 60 specimens was selected to include 30 for which EHV-1 or EHV-4 was detected (15 EHV-1 positive plus 15 EHV-4 positive) and another 30 for which neither EHV-1 nor EHV-4 were detected (Supplemental Table 2). An equine respiratory pathogen other than EHV-1 or EHV-4 was detected in 25 of the latter 30 samples, including EHV-2, EHV-5, equine adenovirus 1 (EAdV1), equine rhinitis A (ERAV), equine rhinitis B (ERBV), equine influenza virus (EIV), *Streptococcus equi* sbsp *equi (S. equi equi)*, *Streptococcus equi* sbsp *zooepidemicus*, or *Mycoplasma felis*. The panel included 8 different specimens (nasal exudate, nasal/pharyngeal swab, whole blood, nasal/pharyngeal wash, tracheal wash, placenta, brain, lung) collected from 21 horse breeds plus a mule and miniature donkey, with ages spanning from an 8-month aborted fetus to 22 year-old adult horse. Specimens originated from 24 states (AR, CT, DE, FL, GA, ID, IL, MA, MD, MN, ND, NH, NJ, NV, NY, OH, OR, PA, SC, TX, VA, VT, WA, WI). To further evaluate diagnostic specificity, 16 donkey and mule samples that previously tested positive for EHV-1 using the assay from Elia and co-workers (47) or a herpesviral species PCR (48) were tested. Thirty specimens used to characterize the equid MC1R assay specificity are listed in Supplemental Table 3.

### Nucleic acid extraction and real-time PCR

Nucleic acid extraction was performed with the MagMAX CORE Nucleic Acid Purification Kit following the simple workflow described by the manufacturer and using the KingFisher Flex (Thermo Fisher Scientific) automated magnetic extractor. Nucleic acid was eluted in 100 μL nuclease-free water. The MS2 plasmid was added to the extraction lysis/binding solution to serve as an exogenous DNA extraction control.

Real-time PCR was optimized and validated using the SensiFAST^TM^ Probe No-ROX Kit (Meridian Bioscience, Memphis, TN) and 7 μL of nucleic acid in a total volume of 25 μL. Real-time PCR was performed on ABI 7500 Fast instruments with cycling conditions of 95°C for 5 minutes followed by 40 cycles of 95°C for 10 seconds and 60°C for 50 seconds, as recommended by the PCR kit manufacturer. The results of the five-plex real-time PCR run were analyzed with 7500 software v2.3 (Life Technologies Corp, Thermo Scientific).

### Analytical assay performance evaluation

Analytical sensitivity was evaluated with ten-fold serial dilutions of EHV-1 and EHV-4 viral isolates and synthetic PACs (Integrated DNA Technologies, Coralville, IA) for each assay target so that the limit of detection could be quantified in copy numbers.

Multiplex assay performance was evaluated with competition experiments performed with 4 replicates each: 1) an EHV-1 isolate was serially diluted in the presence of an EHV-4 isolate at a high concentration, 2) an EHV-4 isolate was serially diluted in the presence of an EHV-1 isolate at a high concentration, and 2) combined EHV-1 and −4 isolates serially diluted. Dilutions were performed prior to extraction and the same nucleic acid elutions were tested side-by-side with the singleplex and multiplex versions of the assays.

Analytical specificity of each assay was evaluated *in silico* with PrimerTree to perform Primer-BLAST, most recently on November 1, 2024 (49).

### Diagnostic assay performance evaluation

Results of the EHV1-4MP assay were compared to results of previously published singleplex EHV-1 and EHV-4 real-time PCR assays (47, 50) and currently used at the Molecular Diagnostics Laboratory at the Animal Health Diagnostic Center to evaluate diagnostic sensitivity and specificity.

### Historical diagnostic testing results

Results of EHV-1, EHV-4, and equine respiratory pathogen panel testing at the Molecular Diagnostics Laboratory at the Animal Health Diagnostic Center from 2011 to December 31, 2024 were downloaded using Tableau Software Server Version: 2022.3.6 (20223.23.0507.0956) (Salesforce Inc, Seattle, WA). The equine respiratory pathogen panel includes real-time PCR tests for EAdV1, equine adenovirus 2 (EAdV2), ERAV, ERBV, EIV, equine arteritis virus (EAV), and *S. equi equi* in addition to EHV-1 and EHV-4. Submissions identified for research purposes were excluded from this analysis. This allowed us to identify a panel of samples known to be positive for other common equine pathogens to determine the diagnostic specificity of our newly developed EHV1-4MP assay.

### Statistics

GraphPad Prism 10.0.0.153 for Windows, GraphPad Software, Boston, Massachusetts USA, www.graphpad.com was used for analysis of standard curve data by linear regression, measuring the difference between slopes of serial dilutions by analysis of covariance, for 2×2 test agreement, and to generate plots.

## RESULTS

### EHV target selection

Glycoprotein B (gB) was selected as an equine herpesvirus real-time PCR assay target because it allows efficient differentiation between EHV-1, EHV-4 and other equine herpesviruses (51, 52). Alignment of EHV-1 full-length gB gene sequences revealed 97.42 to 100% nucleotide identity, with 114 single nucleotide polymorphisms (snps) among the 2,943 nucleotide coding sequence. Alignment of 28 EHV-4 full-length gB gene sequences available in GenBank revealed 99.80 to 100% nucleotide identity, with only 12 snps among the 2,928 nucleotide coding sequence. Alignment of gB sequences from EHV-1 (NC_001491.2) and EHV-4 (NC_001844.1) reference genomes yielded 82.9% nucleotide identity with 500 snps.

The EHV-1 gB assay (EHV1gB444) was designed such that the 3’ end of both primers and probe were anchored on nucleotide positions that differed from EHV-8 and −9. However, this assay still detected EHV-8 and EHV-9 synthetic fragments. Despite additional attempts, we were not able to design a real-time PCR assay in the gB gene that only detected EHV-1 because of the high sequence identity between EHV-1, EHV-8, and EHV-9 (93.2% identical sites, 95.4% pairwise identity). We then used the entire EHV-1 genome sequence for assay design and prioritized exclusion of EHV-8 and EHV-9 sequences. Analysis and comparison of the all EHV-1, −8 and −9 sequences available on public databases led to the design of an assay targeting the EHV-1 genome just upstream the ORF1 coding sequence (EHV1g1775 real-time PCR assay).

### Real-time PCR assay *in silico* specificity assessment

The EHV1g1775 forward and reverse primer pair was evaluated for specificity *in silico* using NCBI nucleotide database. All 162 hits (Supplemental Table 4) were EHV-1 sequences and 160/162 generated the expected amplicon size. No off-target hits were identified.

The EHV1gB444 forward and reverse primer pair was evaluated for specificity *in silico* using NCBI nucleotide database. The EHV-1 gB primers were 100% identical to 143 Equid alphaherpesvirus 1 sequences (Supplemental Table 4) and were predicted to generate an 184 base pair amplicon, as expected. Predicted off-target hits included 8 species of *Schistosoma* and one Marasmarcha lunaedactyla sequence (Supplemental Table 4), all hits used the forward primer at both ends of the amplicon with 2-3 mismatches and generated amplicons ranging from 354 to 3,965 base pairs. To verify assay specificity, another query was performed using the forward primer and probe using PrimerTree. The EHV-1 gB forward primer and probe pair were 100% identical to the same 143 Equid alphaherpesvirus 1 sequences and were predicted to generate a 64 base pair amplicon, as expected; no *Schistosoma* or Marasmarcha hits were identified with this search.

EHV-4 gB primer specificity was assessed *in silico* using the NCBI nucleotide database. The EHV-4 gB forward and reverse primers were 100% identical to 30 Equid alphaherpesvirus 4 sequences (Supplemental Table 4) and were predicted to generate an 82 base pair amplicon, as expected. No off-target hits were identified.

Equid MC1R assay primer specificity was assessed *in silico* using the NCBI nt database. The equid MC1R assay forward and reverse primers match identical sequences from the following 8 equid species and are predicted to generate a 73 base pair amplicon: *Equus asinus, Equus caballus, Equus grevyi, Equus hemionus, Equus kiang, Equus przewalskii, Equus quagga burchellii,* and *Equus zebra* (Supplemental Table 4). Sequences from several other species were identified as possible hits but all have 3 to 7 mismatches with the equid MC1R assay primers and/or probe and the position of many of these mismatches are likely to hinder amplification. These species include tapir (3 mismatches), rhinoceros (5 mismatches), beaver (5 mismatches), dolphin (5 mismatches), squirrel (6 mismatches), lynx (6 mismatches), and pig (7 mismatches) (Supplemental Table 4). Tapirs, rhinoceroses, and horses all belong to the taxonomic order Perissodactyla.

### Analytical assay performance

Performance of each singleplex assay was characterized by testing four replicate serial dilutions of a well-characterized viral isolate or equine control on each of three days. All assays demonstrated acceptable performance based on slope, amplification efficiency, and linearity (Table 2). Intra- and inter-assay variation levels were also evaluated and found to be acceptable.

**Table 2.** Singleplex assay performance on replicate serial dilutions.

| Assay | Slope | Amplification efficiency (%) | Linearity ($R^2$) |
| --- | --- | --- | --- |
| EHV1g1775 | -3.353 to -3.436 | 95.45 to 98.72 | 0.9970 to 0.9996 |
| EHV1gB444 | -3.299 to -3.474 | 94.02 to 101.61 | 0.9981 to 0.9999 |
| EHV4gB | -3.231 to -3.479 | 93.84 to 103.94 | 0.9935 to 0.9999 |
| EqMC1R | -3.215 to -3.477 | 93.91 to 104.53 | 0.9976 to 0.9996 |

Multiplex EHV1-4MP assay performance was compared to singleplex assay performance with 3 experiments: 1) serial dilution of individual EHV isolate, 2) serial dilution of target EHV isolate in the presence of the competing EHV isolate maintained at a high concentration, and 3) serial dilution of combined EHV isolates. After four replicate serial dilutions were generated and extracted, multiplex and singleplex real-time PCR were performed. Assay performance and sensitivity were comparable between singleplex and multiplex formats when an EHV isolate was diluted in the absence or presence of a strong competitor (Figure 1) and when EHV isolates were combined before dilution. The slopes generated from amplification of dilution series did not differ between singleplex and multiplex assays for EHV-1g1775 (p=0.6956), EHV-1 gB (p=0.5922), or EHV-4 gB (p=0.8747).

**Figure 1.**
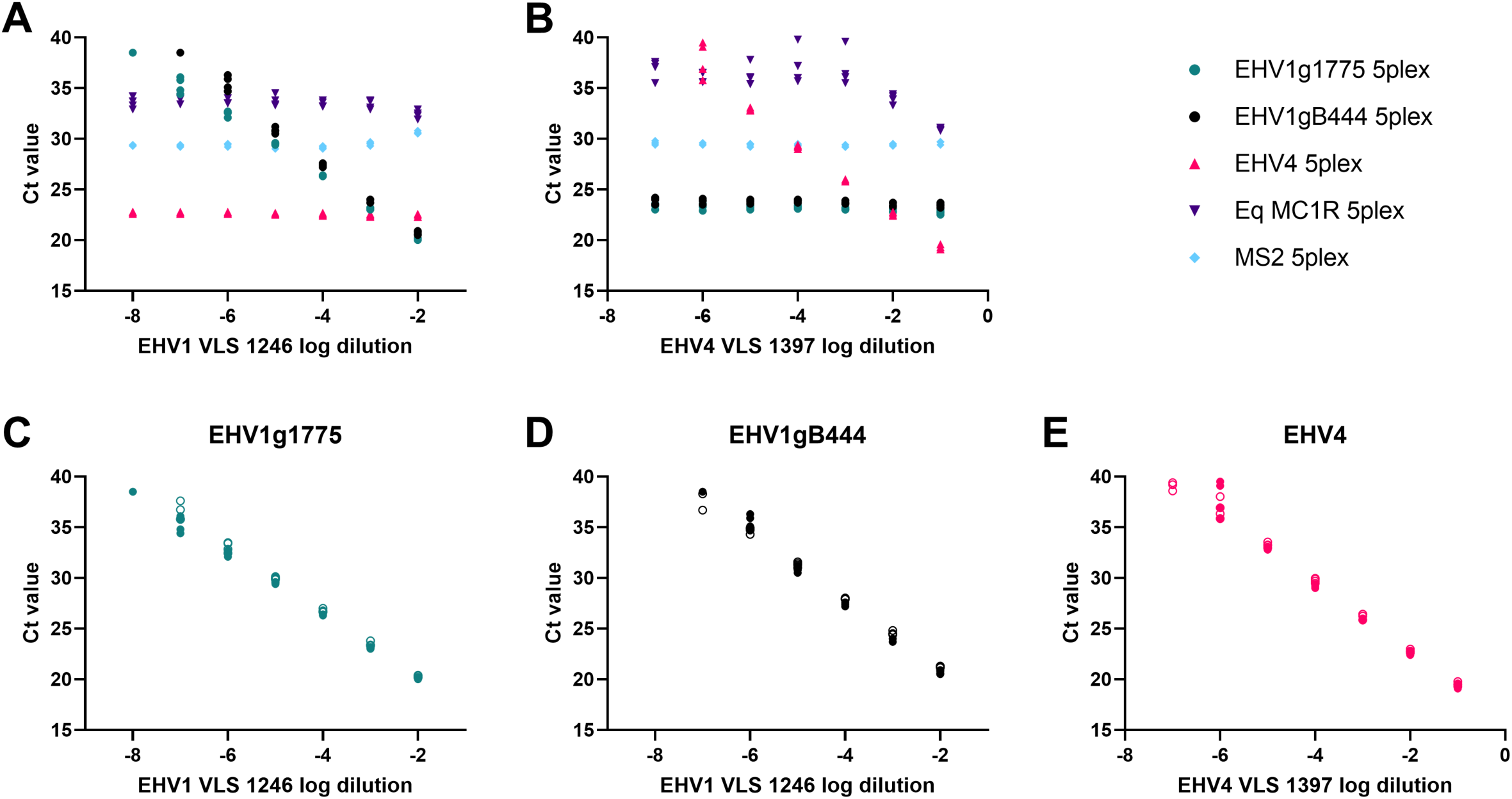
Performance of EHV1-4MP real-time PCR assay in presence of strong competitor. Log dilution of isolate shown on x-axis and Ct value shown on y-axis. A) Serial dilution of EHV-1 isolate in presence of high concentration EHV-4 isolate competitor, B) Serial dilution of EHV-4 isolate in presence of high concentration EHV-1 isolate competitor, C through E) comparison of Ct values determined by multiplex assay (filled symbols) and singleplex assay (open symbols).

### Limit of detection

The limit of detection (LoD) of each assay was initially evaluated from 10-fold serial dilutions of the combined synthetic PACs in 4 replicates. This identified the preliminary LoD to be 10 copies per reaction, based on the lowest copy number for which all replicates were detected. The final LoD was defined by detection of ≥19 of 20 replicates of the combined PACs. The final singleplex LoD was 10 copies per reaction for both EHV-1 assays and the EqMC1R assay, and 15 copies for the EHV-4 assay (Table 3). The final LoD of the multiplex EHV1-4MP real-time PCR assay was 15 copies per reaction (Table 3).

**Table 3.** Limit of detection for EHV1-4MP real-time PCR assay.

| Assay | Singleplex detection |  | Multiplex detection |  |
| --- | --- | --- | --- | --- |
|  | 10 copies | 15 copies | 10 copies | 15 copies |
| EHV1g1775 | 20/20 | not tested | 20/20 | 20/20 |
| EHV1gB444 | 20/20 | not tested | 20/20 | 20/20 |
| EHV4gB | 15/20 | 19/20 | 18/20 | 19/20 |
| EqMC1R | 20/20 | not tested | 20/20 | 20/20 |
| MS2 | not tested | not tested | 20/20 | 20/20 |

### Analytical specificity

Analytical specificity of the EHV1-4MP assay was assessed by performing real-time PCR on a panel of characterized equine respiratory viral (n=16) and bacterial (n=12) isolates. Because EHV-8 and EHV-9 isolates were not available, synthetic DNA fragments were used to evaluate specificity *in vitro*. All EHV-1 isolates were detected by EHV-1 assays, and all EHV-4 isolates were detected by both EHV-4 assays (Table 4). No off-target amplification was detected by the EHV1g1775 (CY5) assay in the EHV1-4MP. The published EHV-1 assay (47) and the EHV1gB444 assay (FAM) in the EHV1-4MP detected EHV-8 and EHV-9 templates, although the EHV1gB444 Ct values were 5-6 cycles later. The previously published EHV-4 assay (50) detected the EHV-3 isolate.

**Table 4.** Analytical specificity of EHV1-4MP assay.

| Pathogen | Isolate name | EHV1-4<br>5plex Ct | EHV-1<br>Ct (47) | EHV-4<br>Ct (50) |
| --- | --- | --- | --- | --- |
| EHV-1 | VLS 1246 | CY5 15.29, FAM 15.48 | 16.74 | Not detected |
| EHV-1 | VLS 727 | CY5 15.12, FAM 15.51 | 16.64 | Not detected |
| EHV-1 clinical isolate | EHV1-035150 | CY5 14.51, FAM 14.62 | 16.60 | Not detected |
| EHV-1 clinical isolate | EHV1-014356 | CY5 22.39, FAM 22.25 | 23.20 | Not detected |
| EHV-2 | VLS 1434 | Not detected | Not detected | Not detected |
| EHV-3 | VLS 1449 | Not detected | Not detected | 36.60 |
| EHV-4 | VLS 1397 | TAMRA 18.99 | Not detected | 17.93 |
| EHV-4 clinical isolate | EHV4-049341 | TAMRA 14.29 | Not detected | 13.18 |
| EHV-4 clinical isolate | EHV4-044856 | TAMRA 16.87 | Not detected | 16.51 |
| EHV-5 | VLS 1462 | Not detected | Not detected | Not detected |
| EHV-8, 10<sup>4</sup> copies | synthetic | FAM 35.62 | 30 | Not detected |
| EHV-9, 10<sup>4</sup> copies | synthetic | FAM 34.04 | 28.32 | Not detected |
| Asinine Herpes isolate | ASHV-010118 | Not detected | Not detected | Not detected |
| Equine Adenovirus 1 | VLS 1535 | Not detected | Not detected | Not detected |
| Equine Arteritis Virus | VLS 1498 | Not detected | Not detected | Not detected |
| Equine Rhinitis A | VLS 1408 | Not detected | Not detected | Not detected |
| Equine Rhinitis B | VLS 1424 | Not detected | Not detected | Not detected |
| Influenza A | VLS 1586 | Not detected | Not detected | Not detected |
| *Actinobacillus equuli* | AEQU-183510 | Not detected | Not detected | Not detected |
| *Actinobacillus lignieresii* | ALIG-080448 | Not detected | Not detected | Not detected |
| *Bordetella bronchiseptica* | BB-088661 | Not detected | Not detected | Not detected |
| *Escherichia coli* | ATCC 25922 | Not detected | Not detected | Not detected |
| *Klebsiella oxytoca* | KO-088499 | Not detected | Not detected | Not detected |
| *Pasteurella caballi* | PCAB-057517 | Not detected | Not detected | Not detected |
| *Pseudomonas aeruginosa* | ATCC 27853 | Not detected | Not detected | Not detected |
| *Rhodococcus equi* | REQU-280087 | Not detected | Not detected | Not detected |
| *Streptococcus dysgalactiae*<br>*sbsp equisimilis* | SDSE-086408 | Not detected | Not detected | Not detected |
| *Streptococcus equi equi* | SEQU-086407 | Not detected | Not detected | Not detected |
| *Streptococcus zooepidemicus* | SZOO-088826 | Not detected | Not detected | Not detected |
| *Taylorella equigenitalis* | NVSL-TE-001 | Not detected | Not detected | Not detected |

### Diagnostic sensitivity and specificity

Diagnostic sensitivity and specificity of the EHV1-4MP assay was evaluated on a panel of 60 residual specimens (30 EHV positive and 30 EHV negative samples) submitted for diagnostic testing at the AHDC. Specimens differed across type, breed, age, sex, location, and presence of other equine respiratory pathogen(s) (Supplemental Table 2). All 30 EHV-1 and EHV-4 positive specimens were detected by the EHV1-4MP and respective singleplex published real-time PCR assays (47, 50) and the other 30 specimens were not detected by any assay, yielding estimates of 100% diagnostic sensitivity and 100% diagnostic specificity (Figures 2A, B). Twenty-five of the samples that did not harbor EHV-1 or EHV-4 were positive for one or more of the following equine pathogens: EHV-2, EHV-5, EAdV1, ERAV, ERBV, EIV, *S. equi equi, S. zooepidemicus,* or *Mycoplasma felis*. The EHV1-4MP assay also provided equid MC1R Ct values from 16.34 to 35.42 (Figure 2C). The exogenous extraction control MS2 DNA was detected in 56 of 60 samples with Ct values 32.75 to 38.77; the other 4 samples had weak or no MS2 amplification but had equid MC1R Ct values ≤ 22.6 (Figure 2C).

**Figure 2.**
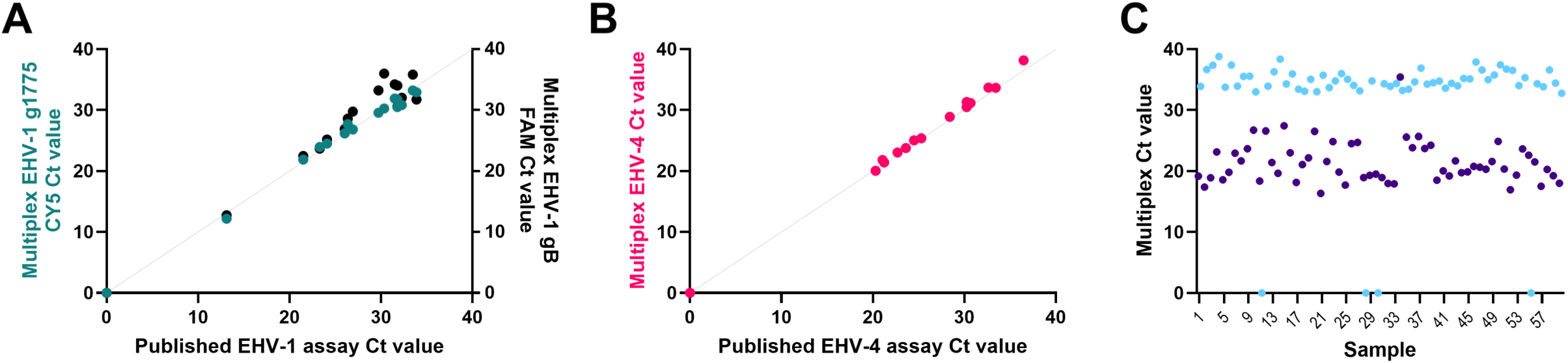
Diagnostic sensitivity and specificity of the EHV1-4MP real-time PCR assay. EHV1-4MP assay results for 60 clinical samples were compared to results of the published singleplex EHV-1 (panel A) and EHV-4 (panel B) assays. EHV-1 Ct values are represented by teal (EHV1g1775) and black (EHV1gB444) circles, and EHV-4 Ct values are represented by pink circles. Not detected results were assigned Ct value of zero. The gray line shows the line of identity. Panel C shows EHV1-4MP control assay results with equid MC1R Ct values represented by purple circles and MS2 DNA exogenous extraction control Ct values represented by light blue circles.

### EHV1-4MP specificity on herpesviruses carried by donkeys

To further characterize the specificity of the EHV1-4MP assay, 16 donkey and mule samples that previously tested positive for EHV-1 using the assay from Elia and co-workers (47) or a herpesviral species PCR (48) were tested (Table 5). Three of the four specimens detected by the published EHV-1 real-time PCR were also detected by the EHV1gB444 assay but not by EHV1g1775, suggesting that these sample may harbor EHV-8 or EHV-9. Sequencing the herpesvirus polymerase gene fragment amplified by the herpesviral species PCR revealed EHV-8 was present in these samples (Table 5). No other herpesviruses carried by these donkeys and mule were detected by the EHV1-4MP assay.

**Table 5.** EHV1-4MP specificity assessment on donkey and mule specimens containing a herpesvirus.

| Sample name | Specimen | EHV1-4MP Ct |  | EHV1 Ct (47) | Herpesvirus pol sequence |
| --- | --- | --- | --- | --- | --- |
|  |  | EHV1g1775 CY5 | EHV1gB444 FAM |  |  |
| ASHV-052072 | Nasal Exudate | ND | 28.90 | 23.11 | EHV-8 |
| ASHV-194585 | Nasal/Pharyngeal swab | ND | 30.98 | 27.77 | EHV-8 |
| ASHV-140854-1 | Nasal Exudate | ND | 32.86 | 26.21 | EHV-8* |
| ASHV-140854-2 | Nasal Exudate | ND | ND | 36.00 | Poor quality sequence |
| ASHV-010118 | Isolate from Nasal Exudate | ND | ND | ND | EHV-7 |
| ASHV-146315 | Nasal/Pharyngeal swab | ND | ND | ND | EHV-7 |
| ASHV-058235 | Fetal membrane | ND | ND | ND | AHV-5 |
| ASHV-142071 | Nasal/Pharyngeal wash | ND | ND | ND | AHV-5 |
| ASHV-185643 | Whole Blood | ND | ND | ND | AHV-5 |
| ASHV-023043-2 | Nasal Exudate | ND | ND | ND | AHV-5 |
| ASHV-023043-3 | Whole Blood | ND | ND | ND | AHV-5 |
| ASHV-229754 | lung, kidney, liver, lymph node | ND | ND | ND | AHV-5 98.61% match |
| ASHV-048022 | Nasal Exudate | ND | ND | ND | AHV-5 98.53% match |
| ASHV-127130 | Nasal Exudate | ND | ND | ND | AHV4-2 |
| ASHV-243608 | Whole Blood | ND | ND | ND | zebra herpesvirus 94.58% match |
| ASHV-073261 | Brain | ND | ND | ND | zebra herpesvirus 93.98% match |

‘ND’ denotes ‘not detected’.
\* EHV-8 sequence identified by next generation sequencing.

### EqMC1R real-time PCR assay specificity

A panel of 30 samples representing a variety of species and specimen types were selected to characterize the equid MC1R real-time PCR assay specificity (Supplemental Table 3). Inclusive equid samples encompassed 10 different horses, 2 different donkeys, a mule, and 3 different zebras. All equid specimens were detected with this assay, including nasal/pharyngeal swabs, whole blood, serum, plasma, cerebrospinal fluid, uterine/vaginal swabs, brain, lung, liver, and formalin-fixed paraffin-embedded liver. For exclusivity testing, non-equid species included cows, goats, sheep, white-tailed deer, reindeer, alpaca, dog, cats, beaver, and dolphin. No non-equid samples were detected by this assay. Exogenous extraction controls were used to confirm successful nucleic acid extraction for all these specimens.

### Historical EHV-1 and EHV-4 detection trends

We investigated the detection rates of EHV-1 and EHV-4 by real-time PCR at the AHDC between 2013 and 2024. EHV-1 was detected in 1.0 to 7.3% of EHV-1 tested samples on an annual basis since 2013 (Figure 3A). EHV-4 detection rates ranged between 2.5 and 7.7% of EHV-4 samples tested since the real-time PCR assay became available at the AHDC in 2015 (Figure 3A). When considered in the context of equine respiratory panel real-time PCR test requests since 2016, EHV-1 was detected in 0.3 to 1.9% of tests annually whereas EHV-4 was detected in 3.9 to 9.4% of tests (Figure 3B). Other equine respiratory pathogens accounted for 16.0 to 30.4% of positive tests, including EAdV1, EAV, ERAV, ERBV, EIV, *S. equi equi, S. zooepidemicus,* and *M. felis,* while approximately 64 to 73% of panel tests did not identify a pathogen (Figure 3B).

**Figure 3.**
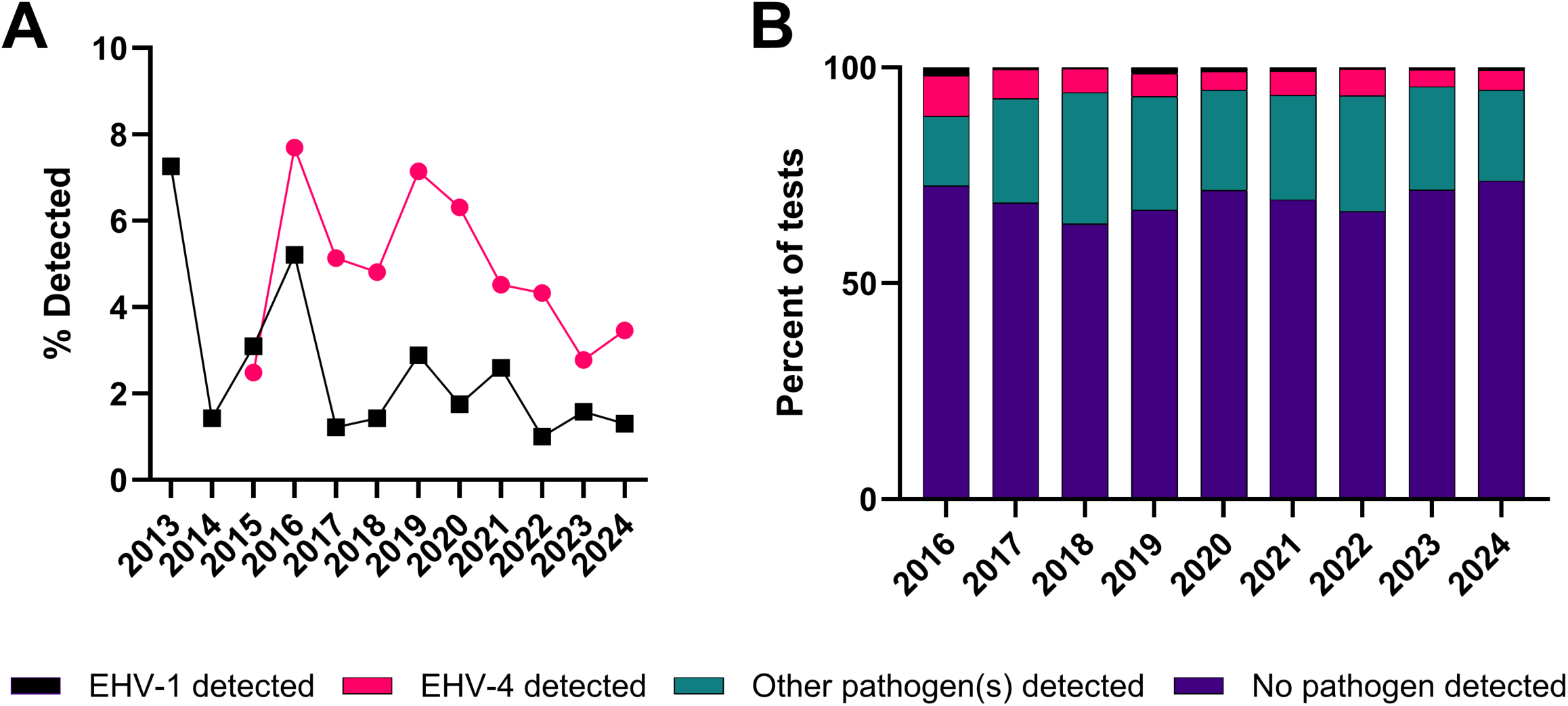
EHV-1 and EHV-4 detected by real-time PCR at the Animal Health Diagnostic Center. A) Percent of EHV-1 and EHV-4 tests with detected result shown on y-axis and year shown on x-axis. EHV-1 detected results are presented in black squares, and EHV-4 results are presented in pink circles. B) Frequency of EHV-1 and EHV-4 detection from equine respiratory pathogen real-time PCR panel test requests. EHV-1 detected results shown in black (0.3 to 1.9%), EHV-4 detected results shown in pink (3.9 to 9.4%), other equine respiratory pathogens detected shown in teal (16.0 to 30.4%), and no pathogen detected shown in purple (63.8 to 73.7%).

This data was further analyzed to assess specimen types that harbor EHV-1 and EHV-4 DNA. EHV-1 was detected in nasal exudate specimens most frequently, followed by blood and respiratory/vaginal swabs; these matrices constitute 85.3% of detected results (Figure 4A). EHV-4 was also detected in nasal exudate specimens predominantly, followed by respiratory swabs; together these 2 matrices comprise 79.9% of detected results (Figure 4B). These results underscore the importance and utility of the EHV1-4 MP assay developed in the present study.

**Figure 4.**
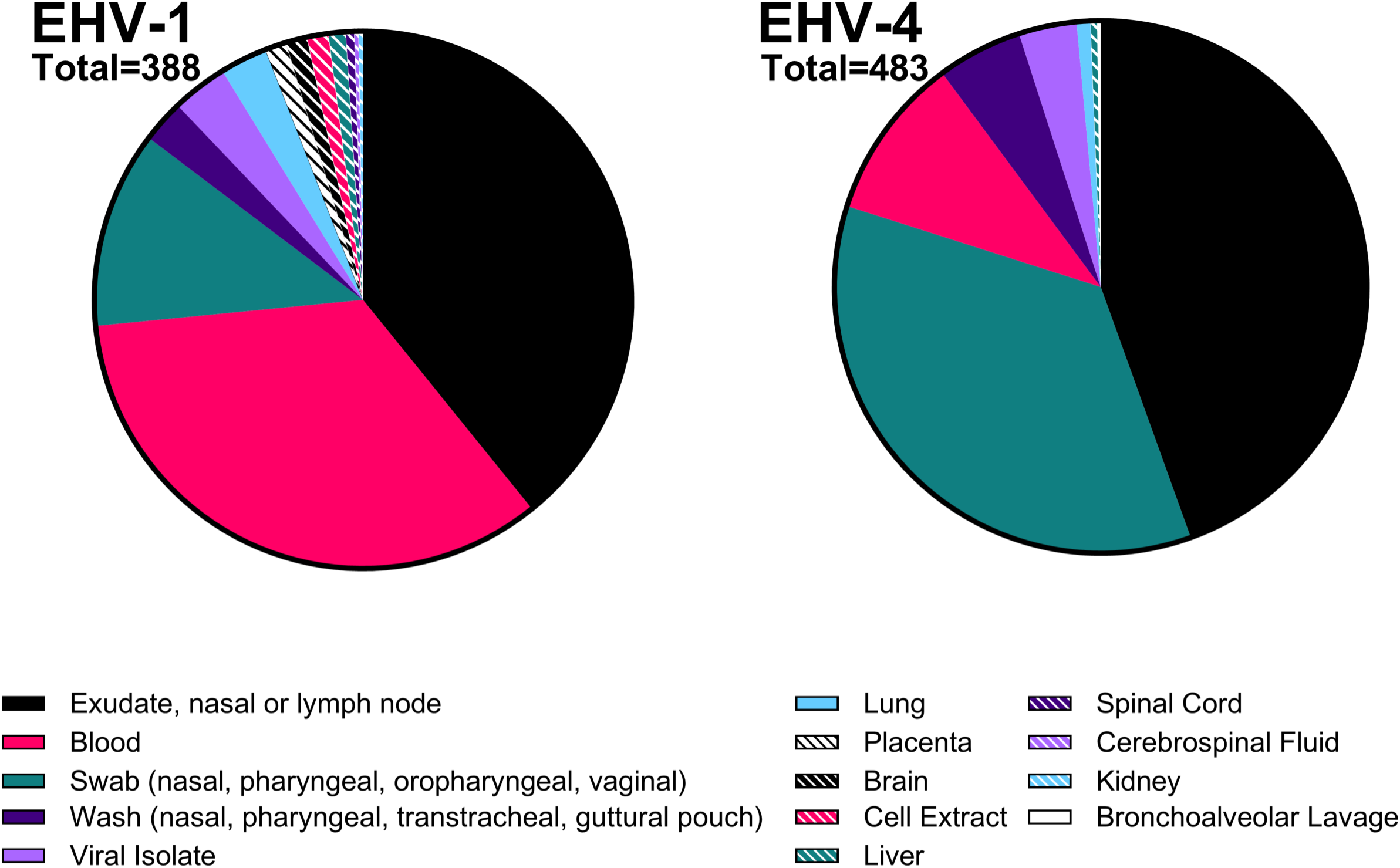
Specimen types of EHV-1 and EHV-4 real-time PCR detected results at the Cornell University Animal Health Diagnostic Center. A) EHV-1 was detected by real-time PCR from 13 different specimen types since 2011. B) EHV-4 was detected by real-time PCR from 8 different specimen types since 2015.

## DISCUSSION

Our goal was to develop a specific and efficient real-time PCR diagnostic assay to enable rapid and simultaneous detection of EHV-1 and EHV-4 infection, in order to support disease control and transmission mitigation efforts. Without prompt and accurate detection, EHV-1 or EHV-4 infections can lead to substantial economic losses due to abortions/neonatal mortality, death following severe disease including EHM, lost training time, and movement restrictions during outbreaks (1, 2, 43). We verified that the multiplex assay format did not affect assay performance and provided sensitive detection of these pathogens (15 copies per reaction). To ensure that this assay provided valid results, both endogenous and exogenous assay controls were included. The endogenous equid control (MC1R) verifies that relevant equine nucleic acid is present in the sample, which may be horse, donkey, mule, hinny, and at least some zebras. The exogenous control provides a measure to assess the presence of PCR inhibitors in the test sample. The importance of monitoring assay performance is illustrated by a real-time PCR study of nasal swab samples in which 27.5% of samples failed quality control (51).

A key feature of the EHV1-4MP assay is its improved specificity over existing assays for EHV-1 detection. The *de novo* EHV1g1775 assay only detects EHV-1, and although the EHV1gB444 assay was designed to exclude amplification of EHV-8 and EHV-9, if high viral loads of these agents are present in the sample, this assay will still cross react and amplify these pathogens. Both EHV-1 assays (EHV1g1775 and EHV1gB444) were kept in the final multiplex assay to increase sensitivity for EHV-1 detection as well as to monitor for EHV-8 or EHV-9, as the prevalence of the latter may be underestimated. Existing EHV-1 gB assays either share 100% identity or differ by 2-3 nucleotides from EHV-8 and EHV-9 gB sequence (47, 50). EHV-1 and EHV-8 sequence similarity contributed to the misdiagnosis of two equine abortions caused by EHV-8 that were initially attributed to EHV-1 (31), and there could be more unrecognized misdiagnoses. Unfortunately, neither EHV-8 nor EHV-9 isolates or characterized positive samples were available for *in vitro* analytical specificity assessment, and so synthetic fragments were utilized for analytical testing. Considering this, and to better characterize the EHV1-4MP assay, we also tested 16 donkey and mule samples that were detected by an EHV-1 real-time PCR (47) or a herpesviral species PCR assay (48). The EHV1g1775 did not detect any samples, indicating that EHV-1 was not present. However, the EHV1gB444 assay detected 3 samples with later Cts than the Elia et al assay (47), suggesting that EHV-8 could be present in these samples. Sequences obtained from these 3 samples confirmed the presence of EHV-8. None of the other donkey samples which were confirmed by sequencing to be positive for EHV-7 or AHV-5 were detected by the EHV1-4MP assay, confirming that it does not amplify these herpesviruses.

A previously described EHV-4 real-time PCR assay (50) amplified EHV-3 DNA in this study, despite only 81% nucleotide identity over the assay target region. Although the differences between EHV-4 and EHV-3 in transmission, tissue distribution, and pathogenesis make misdiagnosis unlikely, it is important to know the inclusivity and exclusivity of assays being used. The *de novo* EHV-4 gB real-time PCR assay validated herein did not detect any other equine pathogens.

Surveillance studies of equine respiratory pathogens by real-time PCR of nasal secretions from thousands of equids with acute fever and/or respiratory symptoms detected EHV-1 in 0.7 – 3.0% of submissions and EHV-4 was detected in 6.6 – 10.8% (6–8). These rates are consistent with those detected by equine respiratory panel real-time PCR tests at the Cornell University Animal Health Diagnostic Center (AHDC) since 2016, where EHV-1 was detected in 0.3 – 1.9% of cases and EHV-4 was detected in 3.9 – 9.4% (Figure 3B). This pattern of higher EHV-4 prevalence than EHV-1 is supported by higher rate of latent EHV-4 detection (83 vs 26%, 41). Other viral and bacterial equine respiratory pathogens including EIV, ERAV, ERBV, EAdV1, EAdV2, EAV, *S. equi,* and *M. felis* were also tested for by real-time PCR at the AHDC equine respiratory panel or by these other studies, diagnosing approximately 20-30% of cases. Notably, these pathogens were not detected in the remaining ∼70% of cases.

In conclusion, we developed a multiplex real-time PCR assay that provides simultaneous, sensitive, and rapid detection of EHV-1 and EHV-4 DNA with improved specificity compared to previously described assays. The EHV1-4MP assay also verifies sample integrity and assay performance at the same time. The multiplex format enables testing for two pathogens and two assay controls within a single well, which may decrease costs and increase workflow efficiency.

## Acknowledgements

We thank Rebecca Franklin-Guild, AHDC Bacteriology Technical Manager, for generously sharing bacterial isolates for this study. This study was funded by the Animal Health Diagnostic Center.

## Data availability statement

The data analyzed in this study are available from the corresponding author.

---

**Supplemental Table 1.**
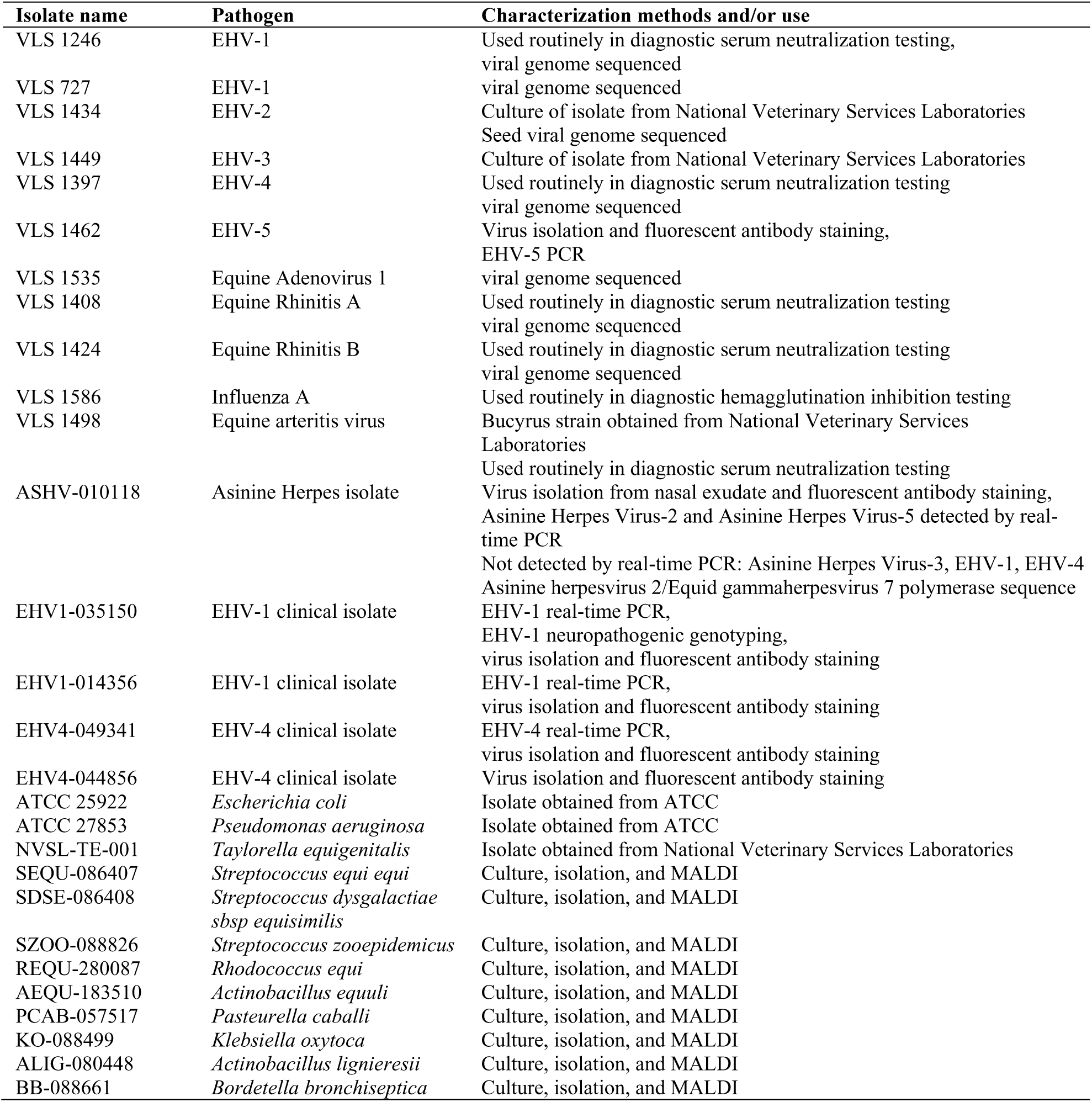
Isolates and clinical specimens used to characterize analytical specificity of the EHV1-4MP assay.

**Supplemental Table 2.** Clinical specimens used for diagnostic sensitivity and specificity estimates.

| Sample ID | Breed<sup>1</sup> | Sex<sup>1</sup> | Age<sup>1</sup> | State<sup>2</sup> | Specimen | Pathogen(s) detected<sup>3</sup> |
| --- | --- | --- | --- | --- | --- | --- |
| 015252 | NS | NS | NS | FL | Nasal Exudate | EHV-1 |
| 017038 | Quarter Horse | Mare | 13 years | VT | Nasal Exudate | EHV-1 |
| 033746 | NS | Castrate | 20 years | MD | Nasal Exudate | EHV-1, *S. equi equi* |
| 038911 | Warmblood | Castrate | 14 years | IL | Nasal Exudate | EHV-1, *S. equi equi* |
| 039219 | Thoroughbred | Castrate | 13 years | CT | Nasal/Pharyngeal Wash | EHV-1 |
| 047621 | Thoroughbred | M | Fetus 10 months | NY | Placenta | EHV-1 |
| 058244 | NS | Mare | NS | MN | Blood, Whole, EDTA | EHV-1 |
| 075126 | Oldenburg | Mare | 1 year | PA | Nasal/Pharyngeal Swab | EHV-1 |
| 076863 | Morgan | Castrate | 16 years | VT | Blood, Whole, EDTA | EHV-1 |
| 095375 | Standardbred | Castrate | 3 years | NY | Brain | EHV-1 |
| 119084 | Standardbred | Mare | Fetus 8 months | NY | Lung | EHV-1 |
| 158668 | Quarter Horse | Stallion | 21 years | VT | Nasal/Pharyngeal Swab | EHV-1 |
| 232536 | Hackney Pony | Mare | 16 years | VA | Swab | EHV-1 |
| 234604 | Thoroughbred | NS | Fetus 8.5 months | NY | Placenta | EHV-1 |
| 280737 | Quarter Horse | Mare | 8 months | ND | Blood, Whole, EDTA | EHV-1 |
| 018436 | Quarter Horse | Mare | NS | TX | Blood, Whole, EDTA | EHV-4 |
| 031926 | Warmblood | Colt | Weanling | NY | Nasal/Pharyngeal Swab | EHV-4 |
| 038649 | Warmblood | Stallion | NS | NY | Nasal/Pharyngeal Swab | EHV-4 |
| 046878 | NS | Mare | 4 years | PA | Nasal/Pharyngeal Swab | EHV-4 |
| 050187 | NS | NS | NS | MA | Nasal Exudate | EHV-4 |
| 059640 | Warmblood | Mare | 1 year | NY | Nasal Exudate | EHV-4, EAdV1 |
| 071577 | Quarter Horse | Castrate | 1 year | OH | Nasal/Pharyngeal Swab | EHV-4, *S. equi equi* |
| 076009 | Haflinger | Mare | 14 years | WA | Blood, Whole, EDTA | EHV-4 |
| 081575 | Arabian | Mare | 6 years | NY | Nasal/Pharyngeal Swab | EHV-4 |
| 177693 | Mule | Mare | 5 years | WA | Swab | EHV-4 |
| 184294 | Quarter Horse | Mare | 2 years | OR | Nasal/Pharyngeal Swab | EHV-4 |
| 191045 | Belgian | Castrate | 4 years | VT | Blood, Whole, EDTA | EHV-4 |
| 220164 | Quarter Horse | Mare | 4 months | MD | Nasal Exudate | EHV-4, EIV |
| 236302 | American Saddle Horse | Stallion | 2 years | NY | Nasal/Pharyngeal Swab | EHV-4 |
| 284915-2 | Thoroughbred | Mare | 6 months | AR | Nasal Exudate | EHV-4 |
| 103495 | Miniature Horse | Stallion | 1 year | TX | Nasal Exudate | EHV-2, ERBV |
| 130518 | Quarter Horse | Stallion | 3 months | TX | Nasal Exudate | EHV-2 |
| 222914 | Quarter Horse | Stallion | 9 months | TX | Nasal Exudate | EHV-2 |
| 211162 | Pony | Mare | 15 years | OH | Nasal Exudate | EHV-5, ERBV |
| 275896 | Thoroughbred | Mare | 20 years | GA | Tracheal wash | EHV-5 |
| 101424 | Thoroughbred | Castrate | 10 years | FL | Nasal/Pharyngeal Wash | EHV-5 |
| 018530 | Haflinger | NS | 10 years | PA | Nasal Exudate | EAdV1 |
| 123882 | Hanoverian | Castrate | 13 years | TX | Nasal/Pharyngeal Swab | EAdV1 |
| 141453 | Quarter Horse | Mare | NS | TX | Nasal Exudate | EAdV1 |
| 020717 | Miniature Horse | Mare | 20 years | MD | Nasal/Pharyngeal Swab | ERAV |
| 133914 | Morgan | Castrate | 5 years | NY | Nasal/Pharyngeal Swab | ERAV |
| 265171 | Warmblood | Castrate | 2 years | PA | Nasal/Pharyngeal Swab | ERAV |
| 025664 | NS | NS | NS | PA | Nasal Exudate | ERBV |
| 011360 | Swedish Warmblood | Castrate | 10 years | DE | Nasal/Pharyngeal Swab | ERBV |
| 020831 | Percheron | Stallion | 1 year | NY | Nasal Exudate | ERBV |
| 017841 | Quarter Horse | Castrate | 22 years | WI | Nasal Exudate | *S. equi equi* |
| 014714 | Thoroughbred cross | Mare | 10 years | OH | Nasal Exudate | *S. equi equi* |
| 009213 | NS | NS | NS | PA | Nasal Exudate | *S. equi equi* |
| 062293 | Quarter Horse | Castrate | 7 years | TX | Nasal/Pharyngeal Wash | *S. zooepidemicus* |
| 228007 | Warmblood | Mare | 4 years | IL | Nasal/Pharyngeal Swab | *S. zooepidemicus* |
| 268893 | Quarter Horse | Stallion | 3 years | ID | Nasal/Pharyngeal Swab | *S. zooepidemicus* |
| 078599 | Miniature donkey | Mare | 4 years | NH | Nasal/Pharyngeal Wash | EIV, *M. felis* |
| 140440 | Mustang | Castrate | 13 years | SC | Nasal/Pharyngeal Swab | EIV |
| 016524 | Quarter Horse | Stallion | 8 years | NV | Nasal Exudate | EIV |
| 234740 | Quarter Horse | Stallion | 3 years | OH | Nasal Exudate | EIV |
| 199895 | Quarter Horse | Castrate | 12 years | VT | Nasal/Pharyngeal Swab | None |
| 284901 | Missouri Fox Trotter | Castrate | 6 years | DE | Nasal Exudate | None |
| 284915-1 | Thoroughbred | Mare | 1 year | AR | Nasal Exudate | None |
| 285653 | Thoroughbred | Mare | 1 year | PA | Nasal Exudate | None |
| 285761 | Paint | Mare | 5 years | NJ | Nasal Exudate | None |

<sup>1</sup> 'NS' indicates not specified
<sup>2</sup> State abbreviations: AR: Arkansas, CT: Connecticut, DE: Delaware, FL: Florida, GA: Georgia, ID: Idaho, IL: Illinois, MA: Massachusetts, MD: Maryland, MN: Minnesota, ND: North Dakota, NH: New Hampshire, NJ: New Jersey, NV: Nevada, NY: New York, OH: Ohio, OR: Oregon, PA: Pennsylvania, SC: South Carolina, TX: Texas, VA: Virginia, VT: Vermont, WA: Washington, WI: Wisconsin
<sup>3</sup> Pathogen abbreviations: EAdV1: equine adenovirus 1, EIV: equine Influenza virus, ERAV: equine Rhinitis A virus, ERBV: equine Rhinitis B virus, *M. felis*: *Mycoplasma felis*, *S. equi equi*: *Streptococcus equi* subspecies *equi*, *S. zooepidemicus*: *Streptococcus zooepidemicus*

**Supplemental Table 3.** Diagnostic specificity assessment of equid MC1R real-time PCR assay.

| Species<br>(common name) | Specimen | Sample identification | Equid MC1R Ct value |
| --- | --- | --- | --- |
| Horse | Nasal/pharyngeal swab | 150241-22-1 | 25.8751 |
| Horse | Nasal/pharyngeal swab | 256704-22-1 | 26.6731 |
| Horse | Nasal exudate | 276046-22-1 | 21.3212 |
| Horse | Brain | 095375-22-1 | 29.8559 |
| Horse | Liver | 004322-23-1 | 29.7414 |
| Horse | Cerebrospinal fluid | 237247-22-1 | 38.6262 |
| Horse | Uterine/vaginal swab | 057423-21-2 | 27.3424 |
| Horse | Serum | 003331-23-1 | 28.0892 |
| Horse | Plasma | 003637-23-1 | 23.3774 |
| Horse | FFPE-fixed liver | 225831-22-1 | 32.0616 |
| Donkey | Nasal/pharyngeal swab | 151194-22-1 | 37.5303 |
| Donkey | Whole blood (EDTA) | 000767-23-1 | 24.2006 |
| Mule | Whole blood (EDTA) | 259967-22-1 | 23.0279 |
| Zebra | Nasal/pharyngeal swab | 275800-22-1 | 34.0127 |
| Zebra | Lung | 173631-22-1 | 23.3725 |
| Zebra | Vaginal swab | 189321-22-1 | 28.2582 |
| Cow | Nasal/pharyngeal swab | 000824-23-1 | Undetermined |
| Cow | Nasal/oropharyngeal<br>swab | 277162-22-1 | Undetermined |
| Goat | Nasal/pharyngeal swab | 193684-22-1 | Undetermined |
| Goat | Nasal/pharyngeal swab | 193684-22-2 | Undetermined |
| Sheep | Whole blood (EDTA) | 001795-23-1 | Undetermined |
| White-tailed deer | Oropharyngeal swab | 206807-22-1 | Undetermined |
| Reindeer | Whole blood (EDTA) | 003060-23-1 | Undetermined |
| Alpaca | Whole blood (EDTA) | 280710-22-31 | Undetermined |
| Dog | Nasal/pharyngeal swab | 004040-23-1 | Undetermined |
| Cat | Nasal/pharyngeal swab | 276225-22-1 | Undetermined |
| Cat | Whole blood (EDTA) | 008140-23-1 | Undetermined |
| Cat | Pleural fluid | 009311-23-1 | Undetermined |
| Beaver | Feces | 080164-20-1 | Undetermined |
| Dolphin | Lymph node | 007298-21-1 | Undetermined |

**Supplemental Table 4.**
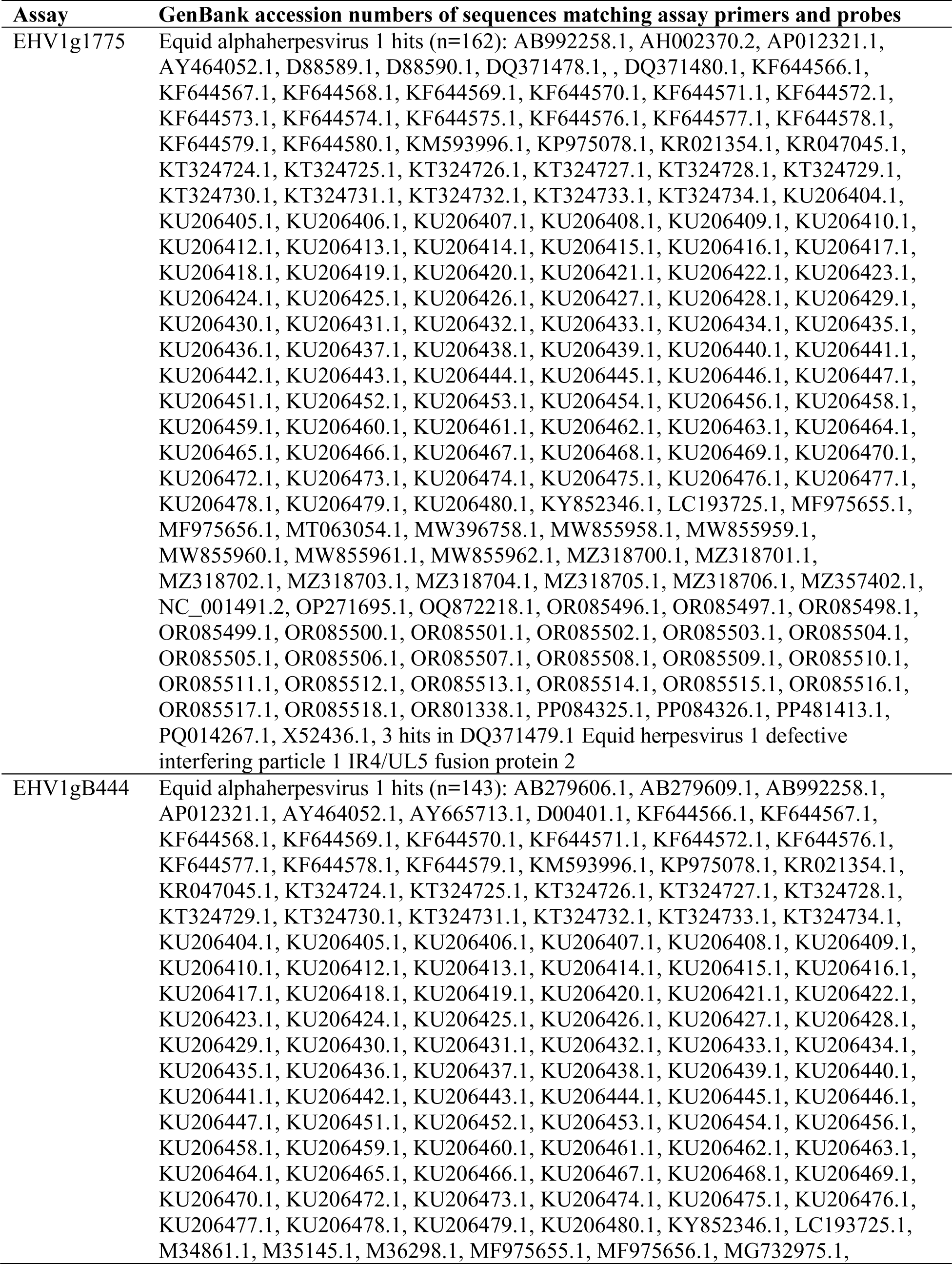
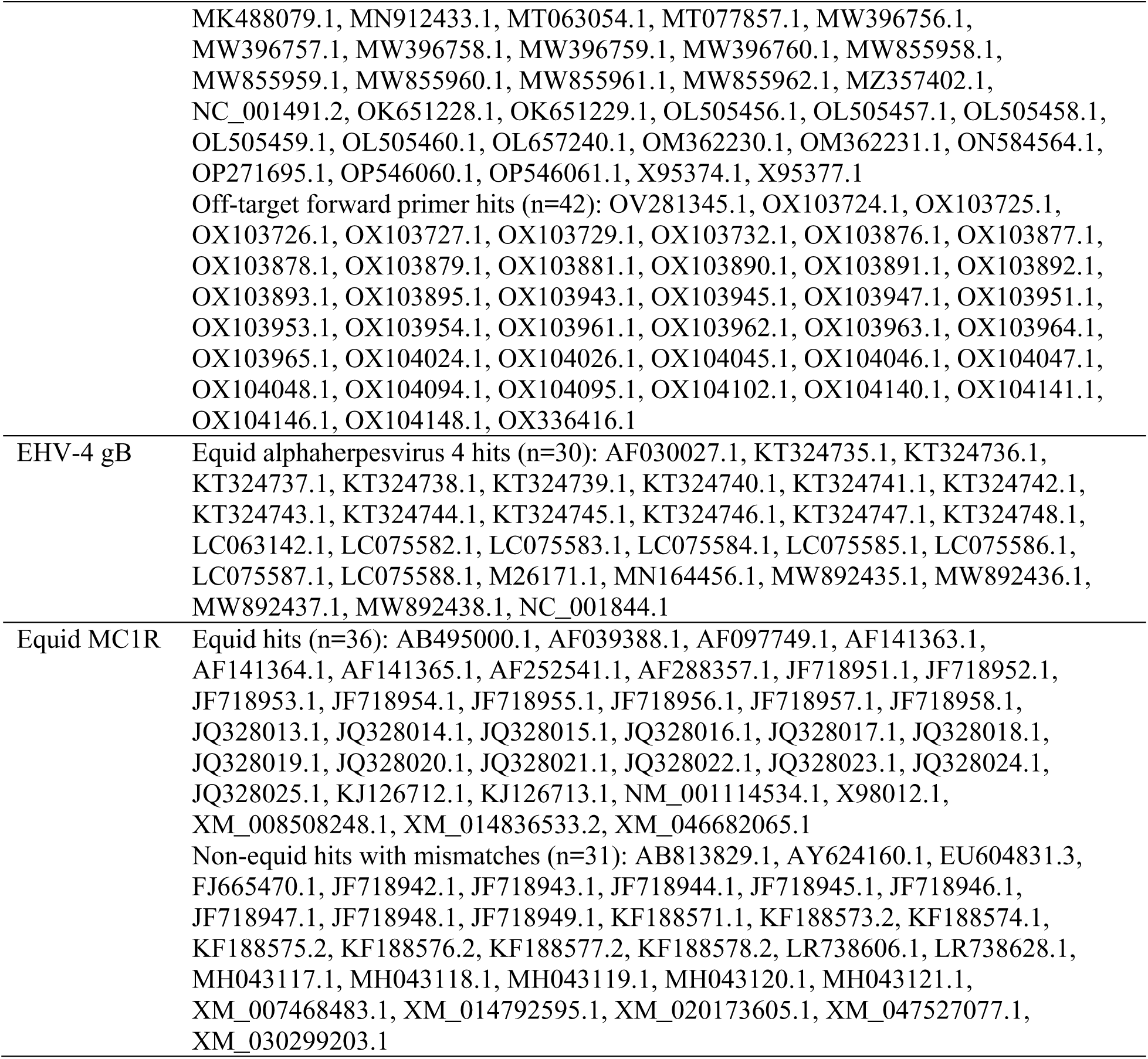
In silico specificity results for EHV1-4MP *de novo* assays.

| **Assay** | **GenBank accession numbers of sequences matching assay primers and probes** |
| --- | --- |
| EHV1g1775 | Equid alphaherpesvirus 1 hits (n=162): AB992258.1, AH002370.2, AP012321.1, AY464052.1, D88589.1, D88590.1, DQ371478.1, , DQ371480.1, KF644566.1, KF644567.1, KF644568.1, KF644569.1, KF644570.1, KF644571.1, KF644572.1, KF644573.1, KF644574.1, KF644575.1, KF644576.1, KF644577.1, KF644578.1, KF644579.1, KF644580.1, KM593996.1, KP975078.1, KR021354.1, KR047045.1, KT324724.1, KT324725.1, KT324726.1, KT324727.1, KT324728.1, KT324729.1, KT324730.1, KT324731.1, KT324732.1, KT324733.1, KT324734.1, KU206404.1, KU206405.1, KU206406.1, KU206407.1, KU206408.1, KU206409.1, KU206410.1, KU206412.1, KU206413.1, KU206414.1, KU206415.1, KU206416.1, KU206417.1, KU206418.1, KU206419.1, KU206420.1, KU206421.1, KU206422.1, KU206423.1, KU206424.1, KU206425.1, KU206426.1, KU206427.1, KU206428.1, KU206429.1, KU206430.1, KU206431.1, KU206432.1, KU206433.1, KU206434.1, KU206435.1, KU206436.1, KU206437.1, KU206438.1, KU206439.1, KU206440.1, KU206441.1, KU206442.1, KU206443.1, KU206444.1, KU206445.1, KU206446.1, KU206447.1, KU206451.1, KU206452.1, KU206453.1, KU206454.1, KU206456.1, KU206458.1, KU206459.1, KU206460.1, KU206461.1, KU206462.1, KU206463.1, KU206464.1, KU206465.1, KU206466.1, KU206467.1, KU206468.1, KU206469.1, KU206470.1, KU206472.1, KU206473.1, KU206474.1, KU206475.1, KU206476.1, KU206477.1, KU206478.1, KU206479.1, KU206480.1, KY852346.1, LC193725.1, MF975655.1, MF975656.1, MT063054.1, MW396758.1, MW855958.1, MW855959.1, MW855960.1, MW855961.1, MW855962.1, MZ318700.1, MZ318701.1, MZ318702.1, MZ318703.1, MZ318704.1, MZ318705.1, MZ318706.1, MZ357402.1, NC_001491.2, OP271695.1, OQ872218.1, OR085496.1, OR085497.1, OR085498.1, OR085499.1, OR085500.1, OR085501.1, OR085502.1, OR085503.1, OR085504.1, OR085505.1, OR085506.1, OR085507.1, OR085508.1, OR085509.1, OR085510.1, OR085511.1, OR085512.1, OR085513.1, OR085514.1, OR085515.1, OR085516.1, OR085517.1, OR085518.1, OR801338.1, PP084325.1, PP084326.1, PP481413.1, PQ014267.1, X52436.1, 3 hits in DQ371479.1 Equid herpesvirus 1 defective interfering particle 1 IR4/UL5 fusion protein 2 |
| EHV1gB444 | Equid alphaherpesvirus 1 hits (n=143): AB279606.1, AB279609.1, AB992258.1, AP012321.1, AY464052.1, AY665713.1, D00401.1, KF644566.1, KF644567.1, KF644568.1, KF644569.1, KF644570.1, KF644571.1, KF644572.1, KF644576.1, KF644577.1, KF644578.1, KF644579.1, KM593996.1, KP975078.1, KR021354.1, KR047045.1, KT324724.1, KT324725.1, KT324726.1, KT324727.1, KT324728.1, KT324729.1, KT324730.1, KT324731.1, KT324732.1, KT324733.1, KT324734.1, KU206404.1, KU206405.1, KU206406.1, KU206407.1, KU206408.1, KU206409.1, KU206410.1, KU206412.1, KU206413.1, KU206414.1, KU206415.1, KU206416.1, KU206417.1, KU206418.1, KU206419.1, KU206420.1, KU206421.1, KU206422.1, KU206423.1, KU206424.1, KU206425.1, KU206426.1, KU206427.1, KU206428.1, KU206429.1, KU206430.1, KU206431.1, KU206432.1, KU206433.1, KU206434.1, KU206435.1, KU206436.1, KU206437.1, KU206438.1, KU206439.1, KU206440.1, KU206441.1, KU206442.1, KU206443.1, KU206444.1, KU206445.1, KU206446.1, KU206447.1, KU206451.1, KU206452.1, KU206453.1, KU206454.1, KU206456.1, KU206458.1, KU206459.1, KU206460.1, KU206461.1, KU206462.1, KU206463.1, KU206464.1, KU206465.1, KU206466.1, KU206467.1, KU206468.1, KU206469.1, KU206470.1, KU206472.1, KU206473.1, KU206474.1, KU206475.1, KU206476.1, KU206477.1, KU206478.1, KU206479.1, KU206480.1, KY852346.1, LC193725.1, M34861.1, M35145.1, M36298.1, MF975655.1, MF975656.1, MG732975.1, |
|  | MK488079.1, MN912433.1, MT063054.1, MT077857.1, MW396756.1, MW396757.1, MW396758.1, MW396759.1, MW396760.1, MW855958.1, MW855959.1, MW855960.1, MW855961.1, MW855962.1, MZ357402.1, NC_001491.2, OK651228.1, OK651229.1, OL505456.1, OL505457.1, OL505458.1, OL505459.1, OL505460.1, OL657240.1, OM362230.1, OM362231.1, ON584564.1, OP271695.1, OP546060.1, OP546061.1, X95374.1, X95377.1<br>Off-target forward primer hits (n=42): OV281345.1, OX103724.1, OX103725.1, OX103726.1, OX103727.1, OX103729.1, OX103732.1, OX103876.1, OX103877.1, OX103878.1, OX103879.1, OX103881.1, OX103890.1, OX103891.1, OX103892.1, OX103893.1, OX103895.1, OX103943.1, OX103945.1, OX103947.1, OX103951.1, OX103953.1, OX103954.1, OX103961.1, OX103962.1, OX103963.1, OX103964.1, OX103965.1, OX104024.1, OX104026.1, OX104045.1, OX104046.1, OX104047.1, OX104048.1, OX104094.1, OX104095.1, OX104102.1, OX104140.1, OX104141.1, OX104146.1, OX104148.1, OX336416.1 |
| EHV-4 gB | Equid alphaherpesvirus 4 hits (n=30): AF030027.1, KT324735.1, KT324736.1, KT324737.1, KT324738.1, KT324739.1, KT324740.1, KT324741.1, KT324742.1, KT324743.1, KT324744.1, KT324745.1, KT324746.1, KT324747.1, KT324748.1, LC063142.1, LC075582.1, LC075583.1, LC075584.1, LC075585.1, LC075586.1, LC075587.1, LC075588.1, M26171.1, MN164456.1, MW892435.1, MW892436.1, MW892437.1, MW892438.1, NC_001844.1 |
| Equid MC1R | Equid hits (n=36): AB495000.1, AF039388.1, AF097749.1, AF141363.1, AF141364.1, AF141365.1, AF252541.1, AF288357.1, JF718951.1, JF718952.1, JF718953.1, JF718954.1, JF718955.1, JF718956.1, JF718957.1, JF718958.1, JQ328013.1, JQ328014.1, JQ328015.1, JQ328016.1, JQ328017.1, JQ328018.1, JQ328019.1, JQ328020.1, JQ328021.1, JQ328022.1, JQ328023.1, JQ328024.1, JQ328025.1, KJ126712.1, KJ126713.1, NM_001114534.1, X98012.1, XM_008508248.1, XM_014836533.2, XM_046682065.1<br>Non-equid hits with mismatches (n=31): AB813829.1, AY624160.1, EU604831.3, FJ665470.1, JF718942.1, JF718943.1, JF718944.1, JF718945.1, JF718946.1, JF718947.1, JF718948.1, JF718949.1, KF188571.1, KF188573.2, KF188574.1, KF188575.2, KF188576.2, KF188577.2, KF188578.2, LR738606.1, LR738628.1, MH043117.1, MH043118.1, MH043119.1, MH043120.1, MH043121.1, XM_007468483.1, XM_014792595.1, XM_020173605.1, XM_047527077.1, XM_030299203.1 |

## REFERENCES

1. Allen G, Kydd J, Slater J, Smith K. 2004. Equid herpesvirus 1 and equid herpesvirus 4 infections., p. 829–859. *In* Infectious Diseases of Livestock.

2. Oladunni FS, Horohov DW, Chambers TM. 2019. EHV-1: A Constant Threat to the Horse Industry. Front Microbiol 10:2668.

3. Pusterla N, Bain F, James K, Mapes S, Kenelty K, Barnett DC, Gaughan E, Craig B, Chappell DE, Vaala W. 2017. Frequency of molecular detection of equine herpesvirus-4 in nasal secretions of 3028 horses with upper airway infection. Veterinary Record 180:593.

4. Pusterla N, James K, Barnum S, Bain F, Barnett DC, Chappell D, Gaughan E, Craig B, Schneider C, Vaala W. 2022. Frequency of Detection and Prevalence Factors Associated with Common Respiratory Pathogens in Equids with Acute Onset of Fever and/or Respiratory Signs (2008– 2021). Pathogens 11:759.

5. Pusterla N, Kass PH, Mapes S, Johnson C, Barnett DC, Vaala W, Gutierrez C, McDaniel R, Whitehead B, Manning J. 2011. Surveillance programme for important equine infectious respiratory pathogens in the USA. Veterinary Record 169:12.

6. Pusterla N, Bain F, James K, Mapes S, Kenelty K, Barnett DC, Gaughan E, Craig B, Chappell DE, Vaala W. 2017. Frequency of molecular detection of equine herpesvirus-4 in nasal secretions of 3028 horses with upper airway infection. Veterinary Record 180:593.

7. Pusterla N, James K, Barnum S, Bain F, Barnett DC, Chappell D, Gaughan E, Craig B, Schneider C, Vaala W. 2022. Frequency of Detection and Prevalence Factors Associated with Common Respiratory Pathogens in Equids with Acute Onset of Fever and/or Respiratory Signs (2008– 2021). Pathogens 11:759.

8. Pusterla N, Kass PH, Mapes S, Johnson C, Barnett DC, Vaala W, Gutierrez C, McDaniel R, Whitehead B, Manning J. 2011. Surveillance programme for important equine infectious respiratory pathogens in the USA. Veterinary Record 169:12.

9. Cantón GJ, Navarro MA, Asin J, Chu P, Henderson EE, Mete A, Uzal FA. 2023. Equine abortion and stillbirth in California: a review of 1,774 cases received at a diagnostic laboratory, 1990–2022. Journal of Veterinary Diagnostic Investigation 35:153–162.

10. Akter R, Legione A, Sansom FM, El-Hage CM, Hartley CA, Gilkerson JR, Devlin JM. 2020. Detection of Coxiella burnetii and equine herpesvirus 1, but not Leptospira spp. or Toxoplasma gondii, in cases of equine abortion in Australia - a 25 year retrospective study. PLoS One 15.

11. Smith KC, Blunden AS, Whitwell KE, Dunn KA, Wales AD. 2003. A survey of equine abortion, stillbirth and neonatal death in the UK from 1988 to 1997. Equine Vet J 35:496–501.

12. Bazanów BA, Frącka AB, Jackulak NA, Staroniewicz ZM, Ploch SM. 2014. A 34-year retrospective study of equine viral abortion in Poland. Pol J Vet Sci 17:607–612.

13. Couroucé A, Normand C, Tessier C, Pomares R, Thévenot J, Marcillaud-Pitel C, Legrand L, Pitel P-H, Pronost S, Lupo C. 2023. Equine Herpesvirus-1 Outbreak During a Show-Jumping Competition: A Clinical and Epidemiological Study. J Equine Vet Sci 128:104869.

14. Walter J, Seeh C, Fey K, Bleul U, Osterrieder N. 2013. Clinical observations and management of a severe equine herpesvirus type 1 outbreak with abortion and encephalomyelitis. Acta Vet Scand 55:19.

15. Pronost S, Legrand L, Pitel PH, Wegge B, Lissens J, Freymuth F, Richard E, Fortier G. 2012. Outbreak of equine herpesvirus myeloencephalopathy in France: a clinical and molecular investigation. Transbound Emerg Dis 59:256–263.

16. Negussie H, Gizaw D, Tessema TS, Nauwynck HJ. 2017. Equine Herpesvirus-1 Myeloencephalopathy, an Emerging Threat of Working Equids in Ethiopia. Transbound Emerg Dis 64:389–397.

17. McFadden AMJ, Hanlon D, McKenzie RK, Gibson I, Bueno IM, Pulford DJ, Orr D, Dunowska M, Stanislawek WL, Spence RP, McDonald WL, Munro G, Mayhew IG. 2016. The first reported outbreak of equine herpesvirus myeloencephalopathy in New Zealand. N Z Vet J 64:125–134.

18. Henninger RW, Reed SM, Saville WJ, Allen GP, Hass GF, Kohn CW, Sofaly C. 2007. Outbreak of Neurologic Disease Caused by Equine Herpesvirus-1 at a University Equestrian Center. J Vet Intern Med 21:157–165.

19. Gryspeerdt A, Vandekerckhove A, Doorselaere J Van, Walle GR Van de, Nauwinck HJ. 2011. Description of an unusually large outbreak of nervous system disorders caused by equine herpesvirus 1 (EHV1) in 2009 in Belgium. Vlaams Diergeneeskd Tijdschr 80.

20. Tong P, Duan R, Palidan N, Deng H, Duan L, Ren M, Song X, Jia C, Tian S, Yang E, Kuang L, Xie J. 2022. Outbreak of neuropathogenic equid herpesvirus 1 causing abortions in Yili horses of Zhaosu, North Xinjiang, China. BMC Vet Res 18.

21. Mureşan A, Mureşan C, Siteavu M, Avram E, Bochynska D, Taulescu M. 2022. An Outbreak of Equine Herpesvirus-4 in an Ecological Donkey Milk Farm in Romania. Vaccines (Basel) 10.

22. Pavulraj S, Eschke K, Theisen J, Westhoff S, Reimers G, Andreotti S, Osterrieder N, Azab W. 2021. Equine Herpesvirus Type 4 (EHV-4) Outbreak in Germany: Virological, Serological, and Molecular Investigations. Pathogens 10.

23. Ryt-Hansen P, Johansen VK, Cuicani MM, Larsen LE, Hansen S. 2024. Outbreak of equine herpesvirus 4 (EHV-4) in Denmark: tracing patient zero and viral characterization. BMC Vet Res 20:1–12.

24. Walker PJ, Siddell SG, Lefkowitz EJ, Mushegian AR, Adriaenssens EM, Alfenas-Zerbini P, Dempsey DM, Dutilh BE, García ML, Curtis Hendrickson R, Junglen S, Krupovic M, Kuhn JH, Lambert AJ, Łobocka M, Oksanen HM, Orton RJ, Robertson DL, Rubino L, Sabanadzovic S, Simmonds P, Smith DB, Suzuki N, Van Doorslaer K, Vandamme AM, Varsani A, Zerbini FM. 2022. Recent changes to virus taxonomy ratified by the International Committee on Taxonomy of Viruses (2022). Arch Virol 167:2429–2440.

25. Vissani MA, Damiani AM, Barrandeguy ME. 2021. Equine Coital Exanthema: New Insights on the Knowledge and Leading Perspectives for Treatment and Prevention. Pathogens 10.

26. Roizman B, Desrosiers R, Fleckenstein B, Lopez C, Minson A, Studdert M. 1995. Herpesviridae, p. 114–127. In Murphy, FA, Fauquet, CM, Bishop, DHL, Ghabrial, SA, Jarvis, AW, Martelli, GP, Mayo, MA, Summers, MD (eds.), Virus Taxonomy. Sixth Report of the International Committee on Taxonomy of Viruses. Springer-Verlag, Vienna.

27. Browning GF, Ficorilli N, Studdert MJ. 1988. Asinine herpesvirus genomes: comparison with those of the equine herpesviruses. Arch Virol 101:183–190.

28. Wang T, Hu L, Wang Y, Liu W, Liu G, Zhu M, Zhang W, Wang C, Ren H, Li L. 2022. Identification of equine herpesvirus 8 in donkey abortion: a case report. Virol J 19.

29. Wang T, Hu L, Liu M, Wang T, Hu X, Li Y, Liu W, Li Y, Wang Y, Ren H, Zhang W, Wang C, Li L. 2022. The Emergence of Viral Encephalitis in Donkeys by Equid Herpesvirus 8 in China. Front Microbiol 13.

30. Liu C, Guo W, Lu G, Xiang W, Wang X. 2012. Complete genomic sequence of an equine herpesvirus type 8 Wh strain isolated from China. J Virol 86:5407–5407.

31. Garvey M, Suárez NM, Kerr K, Hector R, Moloney-Quinn L, Arkins S, Davison AJ, Cullinane A. 2018. Equid herpesvirus 8: Complete genome sequence and association with abortion in mares. PLoS One 13:e0192301.

32. Fukushi H, Tomita T, Taniguchi A, Ochiai Y, Kirisawa R, Matsumura T, Yanai T, Masegi T, Yamaguchi T, Hirai K. 1997. Gazelle herpesvirus 1: a new neurotropic herpesvirus immunologically related to equine herpesvirus 1. Virology 227:34–44.

33. Taniguchi A, Fukushi H, Matsumura T, Yanai T, Masegi T, Hirai K. 2000. Pathogenicity of a new neurotropic equine herpesvirus 9 (gazelle herpesvirus 1) in horses. J Vet Med Sci 62:215–218.

34. Wohlsein P, Lehmbecker A, Spitzbarth I, Algermissen D, Baumgärtner W, Böer M, Kummrow M, Haas L, Grummer B. 2011. Fatal epizootic equine herpesvirus 1 infections in new and unnatural hosts. Vet Microbiol 149:456–460.

35. Tau R, Ferreccio C, Bachir N, Torales F, Romera S, Maidana S. 2023. Comprehensive analysis of equid herpesvirus recombination: an insight into the repeat regions. J Equine Vet Sci 104916.

36. Greenwood AD, Tsangaras K, Ho SYW, Szentiks CA, Nikolin VM, Ma G, Damiani A, East ML, Lawrenz A, Hofer H, Osterrieder N. 2012. A Potentially Fatal Mix of Herpes in Zoos. Current Biology 22:1727–1731.

37. Foote CE, Love DN, Gilkerson JR, Whalley JM. 2004. Detection of EHV-1 and EHV-4 DNA in unweaned Thoroughbred foals from vaccinated mares on a large stud farm. Equine Vet J 36:341– 345.

38. Slater JD, Borchers K, Thackray AM, Field HJ. 1994. The trigeminal ganglion is a location for equine herpesvirus 1 latency and reactivation in the horse. Journal of General Virology 75:2007– 2016.

39. Giessler KS, Samoilowa S, Soboll Hussey G, Kiupel M, Matiasek K, Sledge DG, Liesche F, Schlegel J, Fux R, Goehring LS. 2020. Viral Load and Cell Tropism During Early Latent Equid Herpesvirus 1 Infection Differ Over Time in Lymphoid and Neural Tissue Samples From Experimentally Infected Horses. Front Vet Sci 7:621.

40. Borchers K, Wolfinger U, Ludwig H. 1999. Latency-associated transcripts of equine herpesvirus type 4 in trigeminal ganglia of naturally infected horses. J Gen Virol 80:2165–2171.

41. Pusterla N, Mapes S, David Wilson W. 2012. Prevalence of latent alpha-herpesviruses in Thoroughbred racing horses. Vet J 193:579–582.

42. Carvelli A, Nielsen SS, Paillot R, Broglia A, Kohnle L. 2022. Clinical impact, diagnosis and control of Equine Herpesvirus-1 infection in Europe. EFSA Journal 20:7230.

43. Lunn DP, Burgess BA, Dorman DC, Goehring LS, Gross P, Osterrieder K, Pusterla N, Soboll Hussey G. 2024. Updated ACVIM consensus statement on equine herpesvirus-1. J Vet Intern Med 38:1290–1299.

44. World Organisation for Animal Health. 2024. EQUINE RHINOPNEUMONITIS (INFECTION WITH VARICELLOVIRUS EQUIDALPHA1)Manual of Diagnostic Tests and Vaccines for Terrestrial Animals, 13th ed.

45. Untergasser A, Cutcutache I, Koressaar T, Ye J, Faircloth BC, Remm M, Rozen SG. 2012. Primer3-new capabilities and interfaces. Nucleic Acids Res 40:e115.

46. Dreier J, Störmer M, Kleesiek K. 2005. Use of bacteriophage MS2 as an internal control in viral reverse transcription-PCR assays. J Clin Microbiol 43:4551–4557.

47. Elia G, Decaro N, Martella V, Campolo M, Desario C, Lorusso E, Cirone F, Buonavoglia C. 2006. Detection of equine herpesvirus type 1 by real time PCR. J Virol Methods 133:70–75.

48. Vandevanter DR, Warrener P, Bennett L, Schultz ER, Coulter S, Garber RL, Rose TM. 1996. Detection and analysis of diverse herpesviral species by consensus primer PCR. J Clin Microbiol 34:1666–1671.

49. Cannon M V., Hester J, Shalkhauser A, Chan ER, Logue K, Small ST, Serre D. 2016. In silico assessment of primers for eDNA studies using PrimerTree and application to characterize the biodiversity surrounding the Cuyahoga River. Sci Rep 6:22908.

50. Diallo IS, Hewitson G, Wright LL, Kelly MA, Rodwell BJ, Corney BG. 2007. Multiplex real-time PCR for the detection and differentiation of equid herpesvirus 1 (EHV-1) and equid herpesvirus 4 (EHV-4). Vet Microbiol 123:93–103.

51. Smith FL, Watson JL, Spier SJ, Kilcoyne I, Mapes S, Sonder C, Pusterla N. 2018. Frequency of shedding of respiratory pathogens in horses recently imported to the United States. J Vet Intern Med 32:1436–1441.

